# Frugivore species richness influences dietary specialisation and network properties in Asian wet tropical forests

**DOI:** 10.64898/2026.01.31.703054

**Authors:** Rintu Mandal, Abhishek Gopal, Arpitha Jayanth, Vatcharavee Sriprasertsil, Saniya Chaplod, Himanshu Lad, Aditya Gadkari, Natasha Desai, Rasika Kadam, Anand Osuri, Sartaj Ghuman, Navendu Page, Bee Choo Strange, Vijak Chimchome, Jahnavi Joshi, Rohit Naniwadekar

**Affiliations:** Manipal Academy of Higher Education, Madhav Nagar, Manipal, Karnataka, India, 576104; Nature Conservation Foundation, 1311, Amritha, 12th Main, Vijayanagar First Stage, Mysuru 570017, India; CSIR-Centre for Cellular and Molecular Biology, Uppal Road, Hyderabad, India; Academy of Scientific and Innovative Research (AcSIR), Ghaziabad, India; National Centre for Biological Sciences, Tata Institute of Fundamental Research, Bengaluru 560065, India; Hornbill Research Foundation, Department of Microbiology, Faculty of Science, Mahidol University, Rama 6 Road, Bangkok 10400, Thailand; Conservation Ecology Program, School of Bioresources & Technology, King Mongkut’s University of Technology Thonburi, 49 Soi Tientalay 25, Bangkhuntien-Chaitalay Road, Bangkok 10150, Thailand; Thackeray Wildlife Foundation, Vaibhav Chambers, Bandra, Mumbai 400051, India; Bhavan’s College, Andheri, Mumbai 400058, India; Department of Forest Biology, Faculty of Forestry, Kasetsart University, Thailand

**Keywords:** Community organisation, Frugivory, Hierarchical Modelling of Species Communities, Niche width, Network analysis, Seed dispersal

## Abstract

**Aim:** To examine how variation in frugivore species richness influences dietary specialisation and the organisation of plant-frugivore interaction networks in tropical forests.

**Location:** Six undisturbed lowland wet tropical forest sites across four biodiversity hotspots in south and south-east Asia.

**Time period:** 2016–2024.

**Major taxa studied:** Avian frugivores and fleshy-fruited woody plants.

**Methods:** We recorded plant-avian frugivore interactions across six undisturbed evergreen forest sites spanning a seven-fold gradient in frugivore species richness, while holding forest type and phylogenetic composition broadly comparable. Using over 4,200 hours of focal observations on 551 fruiting plants, we recorded more than 34,000 feeding visits by 138 frugivore species on 133 plant species. We used a) Joint species distribution models to determine the relative influence of fruit and seed traits, and b) network analyses to evaluate how dietary breadth and network properties varied with frugivore species richness.

**Results:** Across sites, frugivore visitation was primarily explained by fruit and seed morphology, with seed size accounting for an average of 39.7% of explained variation, followed by fruit width (24.4%), fruit crop size (21.9%), and pulp lipid content (14.1%). Frugivores in species-rich communities exhibited narrower dietary breadth (Pearson’s *r* = −0.87 between normalised degree and species richness). Correspondingly, plant-frugivore networks became less connected and nested, and more modular, with increasing frugivore richness (Pearson’s *r* = −0.9, −0.98, and 0.84, respectively).

**Main conclusions:** Increasing frugivore species richness intensifies dietary specialisation, which in turn drives changes in plant-frugivore network structure. These findings highlight how local species richness shapes interaction networks through changes in consumer niche breadth, with implications for the organisation of tropical forest mutualistic communities.

## INTRODUCTION

How species diversity is maintained within ecological communities remains a central question in ecology. Beyond the role of abiotic factors, biotic interactions such as competition play a key role in structuring local communities (HilleRisLambers *et al*., 2012). As richness in the local species pool increases, inter-specific competition for resources is expected to intensify (Gainsbury & Meiri, 2017). Following the modern theory of coexistence (Chesson, 2008), coexistence under increased competition arises through niche specialisation, which reduces niche overlap and facilitates partitioning of resources. In the context of trophic interactions, increasing niche specialisation (e.g., frugivores visiting fruiting plants) can scale up to influence properties of ecological networks, such as connectance (proportion of realised interactions), nestedness (extent to which specialists interact with generalist partners), and modularity (degree to which interactions are compartmentalised into distinct groups) (Schleuning *et al*., 2011). These properties are important to study as they determine community organisation and stability: greater compartmentalisation and specialisation may increase vulnerability to partner loss (Olesen *et al*., 2007; Aizen *et al*., 2012), while nestedness may buffer against extinction cascades by diffusing impacts across the network (Memmott *et al*., 2004). Understanding how niche specialisation and network organisation vary across species richness gradients is therefore vital for predicting the resilience of ecological communities.

Theoretical models predict that increasing richness promotes nestedness and reduces modularity, thereby minimising interspecific competition and allowing multiple species to coexist (Bastolla *et al*., 2009). Yet empirical studies report contrasting patterns: reduced nestedness and greater modularity with higher species richness (Olesen *et al*., 2007; Almeida & Mikich, 2018). These discrepancies in network properties may arise because the consequences of richness depend on how niche widths change. Narrower niches with increasing richness may increase modularity and specialisation but reduce connectance and nestedness, promoting species coexistence by reducing competition (sensu Bastolla et al., 2009). On the other hand, increasing species richness may not alter (Novosolov *et al*., 2018) or even positively influence niche widths (Büchi & Vuilleumier, 2014), which can result in more nested, less modular and specialised communities. Despite theoretical work, empirical data linking species richness, dietary specialisation, and community organisation remains relatively scarce.

Among the various dimensions of niche differentiation, diet is a particularly important one, along which consumers can respond to increasing competition. In species-rich communities, consumers may coexist by narrowing their dietary breadth, a species-level response that is often quantified as greater dietary specialisation (Pianka, 1974). As species interact with a smaller subset of available resources, these species-level responses scale up to influence the organisation of ecological networks. Specifically, reduced dietary breadth can lead to fewer realised interactions per species, resulting in lower connectance and nestedness, while promoting greater network specialisation and modularity (Blüthgen *et al*., 2006; Rezende *et al*., 2009). Although dietary specialisation has been shown to influence network structure (Schleuning *et al*., 2011), few studies have explicitly tested whether increasing species richness drives dietary specialisation and, in turn, alters community organisation (but see Gómez-Martínez *et al*., 2022).

Plant-frugivore systems provide an excellent model to examine these dynamics. Fruits constitute a major food resource in wet tropical forests, often comprising a substantial proportion of the diet of frugivorous birds and mammals (Jordano, 2000). At the ecosystem level, annual fruit production can reach up to 1,000 kg per ha (Jordano, 2000). Plant-frugivore interactions are critical for seed dispersal and the maintenance of plant diversity. However, visitation to fruiting plants is influenced by multiple factors, including morphological trait matching between fruits and frugivores (Bender *et al*., 2018; Naniwadekar *et al*., 2019), fruit availability and nutrient composition (Herrera *et al*., 2011; Pizo *et al*., 2021), and interactions among frugivore species (French & Smith, 2005). Optimal foraging theory predicts that frugivores should preferentially consume fruits they can handle efficiently and that offer high or complementary (i.e., a mix of lipids and carbohydrates) rewards (Blendinger *et al*., 2022). Accordingly, frugivores may exhibit morphological matching with fruit traits, track fruit resource abundance, and/or select fruits with particular nutrient contents. Yet most studies examine these correlates in isolation (but see Rojas *et al*., 2021), and we still know little about their relative contributions, especially under varying levels of community complexity. Local frugivore species richness has been widely used as an integrative proxy of interspecific competition in systems where consumers rely on shared resources (Gainsbury & Meiri, 2017; Freeman *et al*., 2024). In such contexts, higher richness increases the likelihood of niche overlap and competitive interactions. This framework is particularly important for frugivores, which are known to exhibit dietary plasticity in the absence of competitors but may experience stronger constraints on resource use as community richness increases (Naniwadekar *et al*., 2021).

Species-richness gradients provide opportunities to investigate how increasing competition, as a potential consequence of increasing species richness, influences dietary niches and community organisation. However, such gradients are often confounded by underlying environmental gradients, making it difficult to disentangle the effects of abiotic conditions from interspecific competition. “Species richness anomalies” (Swenson *et al*., 2016), where assemblages differ in richness under similar environmental conditions due to historical processes such as dispersal or in-situ speciation, offer a natural experiment to isolate the effects of biotic interactions. Controlling for environmental factors is crucial, as network specialisation has been shown to decrease with elevation and latitude (Dalsgaard *et al*., 2017; Angulo-Ortiz *et al*., 2024). Although several studies have examined relationships between species richness and network properties along environmental gradients (Almeida & Mikich, 2018), evidence from sites with comparable environments but contrasting frugivore richness remains scarce. While sampling across species richness gradients, it is also critical to sample phylogenetically-related communities, as evolutionary relationships between frugivores and fruit species may differ drastically, with consequences for contemporary community organisation. For example, Afrotropical networks, which are phylogenetically distinct from the Neotropical ones, were found to be less specialised, as Afrotropical birds exhibited greater dietary overlap (Dugger *et al*., 2019). Methodological consistency is important too, as differences in sampling approaches can strongly influence the observed relationships between richness and network properties (Brimacombe *et al*., 2023). To draw robust conclusions on how species richness influences the structure and organisation of interaction networks, it is essential to apply uniform sampling methods across sites that differ primarily in species richness.

Given this background, we examine plant-frugivore interactions across six wet tropical sites in south and south-east Asia that span a gradient in frugivore species richness. These sites experience similar climatic conditions and harbour phylogenetically nested assemblages of frugivore and fruiting plant species, allowing meaningful comparisons across richness levels while minimising environmental and historical confounds (Naniwadekar *et al*., 2025).

Specifically, we address the following questions: 1) Does frugivore species richness modify the relative influence of fruit and seed morphology, pulp nutrient content, and fruit crop size on the occurrence of frugivores on fruiting plants? 2) Do frugivores exhibit greater dietary specialisation in communities with higher frugivore species richness? 3) Do plant-frugivore networks in species-rich communities become more modular and specialised, with lower connectance and nestedness? The first question examines whether greater interspecific competition, reflected in higher frugivore species richness, can alter the relative importance of fruit and seed traits, fruit availability, and fruit quality in driving frugivore visits on fruiting plants. Specifically, while fruit and seed morphology impose strong mechanical constraints on frugivore-plant interactions, increased competition may enhance the role of other dimensions such as quantity (fruit availability) and quality (nutrient content) of resources. The second and third questions investigate whether increased competition promotes dietary specialisation, leading to more compartmentalised and specialised communities. We expect that the morphological traits of seeds and fruits will have the strongest influence on frugivore visitations, regardless of frugivore species richness, as they impose mechanical constraints on avian frugivores that prevent interaction. However, in line with the evidence that more diverse communities tend to exhibit greater niche differentiation (MacArthur & Levins, 1967), we expected to find greater dietary specialisation with increasing frugivore species richness, resulting in less connected and nested, and more specialised and modular plant-frugivore communities.

## MATERIALS AND METHODS

### Study area

We conducted this study across six Protected Area sites in south and south-east Asia – five in India and one in Thailand – spanning four biodiversity hotspots: the Western Ghats–Sri Lanka, the Himalaya, the Indo-Burma, and the Sundaland hotspots. These sites experience minimal anthropogenic disturbance to fleshy-fruited plants and frugivorous birds. Among the Indian sites, two were located in the Eastern Himalaya–Namdapha Tiger Reserve (hereafter Namdapha; 27°23′–27°39′N and 96°15′–96°58′E) and Pakke Tiger Reserve (hereafter Pakke; 26°54–27°16′N and 92°36′–93°09′E), one in the Western Ghats–Anamalai Rainforest Research Station of the Anamalai Tiger Reserve (hereafter Anamalai; 10°12′–10°35′N and 76°49′–77°24′E), and two were oceanic island sites in the Andaman Archipelago–South Andaman (hereafter Andaman; 11°07′–12°15′N and 92°30′–92°50′E) and Narcondam (13°30′N and 94°38′E). The Thailand site was located in the Hala-Bala Wildlife Sanctuary (hereafter Bala; 5°44′–5°57′N and 101°46′–101°51′E) (Table 1). According to Whitmore’s (1984) classification of tropical rain forests, all sites can be categorised as tropical lowland evergreen rainforests, as each lies below 1,200 m above mean sea level and receives over 2,000 mm of mean annual rainfall (Fig. 1). There was nearly a sevenfold difference in the number of avian frugivore species documented in this study across sites, ranging from seven species on Narcondam Island (the most species-poor site) to over 40 species in the Eastern Himalayan and Sundaland sites. Further details on the study area are provided in Naniwadekar *et al*., (2025).

**Table 1.** Summary of the data collected across six study sites. The number of avian frugivore species recorded during the sampling period increases from top to bottom. *While the area of the Protected Area is reported here, sampling was carried out in a smaller portion of the park that had lowland tropical evergreen forests.

| Study sites | Area of the Protected Area* | Sampling period | Number of frugivore species recorded | Number of plant species observed | Number of focal plant individuals | Number of frugivore visits | Number of unique interactions | Total focal watch duration (h) |
| --- | --- | --- | --- | --- | --- | --- | --- | --- |
| Narcondam | 6.8 km<sup>2</sup> | Dec' 2019 to Feb' 2020 | 7 | 13 | 41 | 696 | 28 | 246 |
| Andaman | 136 km<sup>2</sup> | Dec' 2022 to Mar' 2024 | 24 | 24 | 93 | 8667 | 170 | 558 |
| Anamalai | 987 km<sup>2</sup> | Nov' 2023 to May 2024 | 26 | 28 | 119 | 3498 | 186 | 714 |
| Namdapha | 1985 km<sup>2</sup> | Nov' 2022 to Apr' 2023 | 40 | 28 | 67 | 5911 | 205 | 402 |
| Bala | 111.5 km<sup>2</sup> | Aug' 2022 to May 2023 | 45 | 22 | 47 | 1423 | 253 | 297 |
| Pakke | 861.95 km<sup>2</sup> | Oct' 2016 to Feb' | 48 | 46 | 184 | 14398 | 326 | 2050 |
|  |  | 2018 |  |  |  |  |  |  |

**Figure 1.**
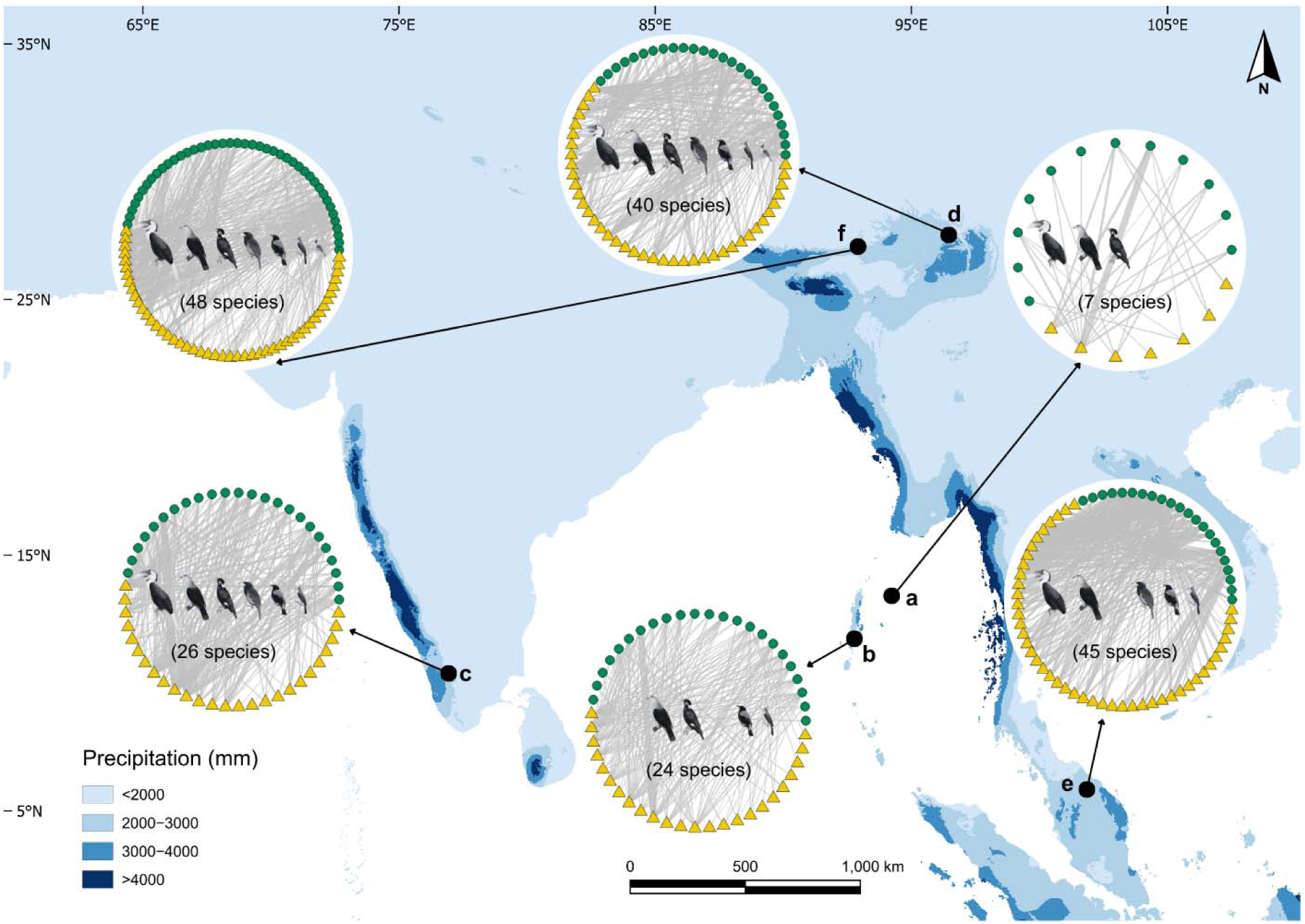
Map of south and south-east Asia with mean annual precipitation (mm) data overlaid in blue shades and our study sites marked in red dots (a - Narcondam, b - Andaman, c - Anamalai, d - Namdapha, e - Bala, and f - Pakke). The large circles represent weighted plant-frugivore bipartite networks for our respective study sites. Within each bipartite network, the green circles represent fruiting plant species, the yellow triangles represent avian frugivore species, and the grey lines represent links between avian frugivores and fruiting plants. Bird illustrations within each large circle represent major frugivore families (left to right: Bucerotidae, Columbidae, Sturnidae, Megalaimidae, Irenidae, Pycnonotidae, and others) in that particular site. The numbers at the bottom of each large circle represent the site-level richness of avian frugivores recorded during the study. Bird illustrations (modified) are taken from https://www.birdsoftheworld.org/, and mean annual precipitation information from WorldClim v2.1 (https://www.worldclim.org/).

### Plant-frugivore interactions

We conducted focal watches on fleshy-fruited woody plants across six sites to record interactions between avian frugivores and fruiting plants (Table 1). Data were collected between 2016 and 2024 during the non-monsoon months (mostly between October and May), when fruit availability is relatively low (Datta, 2001), and interspecific competition for fruits is expected to be higher. We located fruiting plants by surveying existing trails and opportunistic walks in the study areas. We observed fruiting plants bearing ripe fruits, ensuring representation of small, medium and large-seeded plants. Additional details of fruiting trees observed in each site are provided in Table 1 and Naniwadekar *et al*., (2025).

We observed each focal plant for six hours starting at sunrise, except in Pakke, where plants were observed for a full day (see Naniwadekar *et al*., 2019 for details). Each individual plant was observed only once to ensure independence of samples. The duration of focal watches did not affect the number of frugivore species recorded (Fig. S1 in Naniwadekar *et al*., 2025). Two trained observers made observations using 8 × 40 or 8 × 42 binoculars, positioned at a suitable distance to minimise disturbance. We recorded the species identity and number of individual birds visiting the plant. Only birds observed feeding (swallowing or pulp pecking) on fruits were classified as frugivores, and all their subsequent observations were considered as interactions following previous studies (Schleuning *et al*., 2012; Naniwadekar *et al*., 2019). We estimated the ripe fruit crop size of each focal plant following Naniwadekar *et al*., (2019). Avian frugivore richness and observed fruiting plant species were not correlated with each other (Table S2 in Naniwadekar *et al*., 2025), suggesting that differences in the number of observed plant species did not influence the number of frugivore species observed. We conducted a total of 4,267 hours of focal watches on 551 bird-dispersed fleshy-fruited individual woody plants. Further details of focal watches, including site-wise sampling effort, number of species, and individuals observed, are provided in Table 1.

### Bird and fruit traits

We estimated mean fruit and seed width for focal plant species using a digital vernier calliper, measuring at least five fresh fruits and seeds per species. We relied on existing floras and expert knowledge (Navendu Page) for classifying species across different seed size categories for which seed size data was not collected in the field. We used fruit width on a continuous scale, whereas we classified plants as small-seeded (seed width < 5 mm), medium-seeded (5–15 mm), and large-seeded (> 15 mm) following Naniwadekar *et al*., (2019). We used the percentage lipid content by dry weight as a measure of macronutrient content of fruit pulp. We did not use information for non-structural carbohydrates, as comparable data were unavailable for many species or genera, and lipid content and non-structural carbohydrates are reported to be negatively correlated in fleshy fruits (Pizo et al., 2021). We estimated lipid content for 18 of the 28 focal plant species in Anamalai (Table S2), as facilities for fruit processing and storage were available at the Anamalai Rainforest Research Station. For these species, we collected fresh ripe fruits from under focal plants, deseeded them, and oven-dried the pulp at 60°C for 72 hours. The dried pulp samples were analysed at the Kerala Agricultural University, Thrissur, Kerala, following AOAC protocols (20th Edition, 2016, 920.85). Estimation was not possible for fruits with minimal pulp due to the difficulty in obtaining sufficient dry mass for analysis. For species whose fruits could not be analysed directly, we collated percentage lipid content from published literature at the most precise taxonomic level (species/genus/family). Following this, we extracted values for the 83 of the 115 remaining plant species from published literature (Table S2). Lipid content is a phylogenetically conserved trait (Pizo et al., 2021); therefore, the use of secondary data for nutrients is unlikely to significantly impact the interpretation of the results. Moreover, we classified the plants into three pulp lipid categories: low (<10%), medium (10-33%), and high (>33%), following Pizo *et al*., (2021).

We compiled bird functional traits from global trait databases: beak width and hand-wing index (HWI) from AVONET (Tobias *et al*., 2022), and the frugivory degree (percentage fruit in diet) from EltonTraits (Wilman *et al*., 2014). Beak width and HWI represent the frugivore’s capacity for fruit handling and potential for movement for tracking fruit resources or dispersing seeds, respectively (Dehling *et al*., 2014), and the frugivory degree reflects their dietary specialisation on fruits (Pizo et al., 2021). Bird and plant species taxonomy were harmonised by following Clement’s Checklist of Birds and Plants of the World online, respectively.

### Analyses

#### Frugivore occurrence on fruiting plants

To determine the relative importance of different fruit traits and availability in influencing the avian frugivore visitation to fleshy-fruited woody plants, we used the Hierarchical Modelling of Species Communities (HMSC) framework developed by (Ovaskainen & Abrego, 2020). HMSC is a class of joint species distribution models that determines species responses to predictors while accounting for their functional traits and phylogenetic relationships within the community. For each site, we converted the plant-frugivore visitation matrix (number of bird visits per hour to individual focal plants) into a presence-absence matrix. We then modelled the occurrence of each frugivore species on individual fruiting plants as a function of fixed effects, which included seed size, fruit width, pulp lipid content, and the log-transformed ripe fruit crop size. Each focal plant represented one sampling unit. The analysis was conducted separately for five of the six study sites. Bala was excluded from this analysis as fruit morphology data were unavailable for several plant species. Rare frugivores, i.e., avian frugivore species that occurred on less than five percent of the total individual fruiting plants watched in a particular site, were removed from the analysis. It is recommended to drop rare species from the analysis as they interfere with MCMC model convergence (Ovaskainen & Abrego, 2020). Thus, we conducted HMSC analyses for 15 frugivore species each for Anamalai and Andaman, 28 for Namdapha, five for Narcondam, and 20 for Pakke. We included three avian functional traits–beak width, HWI, and frugivory degree into the HMSC framework. To account for phylogenetic effects, we used site-specific time-calibrated bird phylogenetic trees obtained from https://www.birdtree.org/ (Jetz *et al*., 2012). Each site-specific tree was pruned to only include bird species recorded in that site. For species not represented in the phylogeny, data were substituted from their closest congeneric relatives.

We used the R package ‘Hmsc’ to analyse how frugivore occurrences on fruiting trees were influenced by the predictor variables, assuming default prior distributions (Tikhonov *et al*., 2020). We implemented four Markov Chain Monte Carlo (MCMC) chains to sample from the posterior distribution. For each study site, each chain was run for 375,000 iterations, with the first 125,000 iterations discarded as burn-in. The remaining samples were thinned by 1,000, resulting in 250 posterior samples per chain and a total of 1,000 posterior samples across chains. We assessed MCMC convergence using the potential scale reduction factor (PSRF) values, considering chains to have converged when PSRF values < 1.05 (Ovaskainen & Abrego, 2020). We used ‘probit’ models for the presence-absence data and examined model fit using the species-specific Tjur’s *R^2^* values. For each site, we quantified the percentage of variation explained by each fixed effect in the model (seed size, fruit width, pulp lipid content, and log-transformed fruit crop size). We computed the mean percentage variation explained by each fixed effect across the five sites. To examine whether the percentage variance explained differed significantly across predictors, we performed beta regression followed by post-hoc pairwise comparisons using the R packages ‘betareg’ and ‘emmeans’, respectively.

### Frugivore dietary specialisation

To examine the dietary specialisation of avian frugivores across our study sites, we calculated the Normalised Degree (ND) and Blüthgen’s *d*′ indices for each frugivore species using the R-package ‘bipartite’ (Dormann *et al*., 2009). Both indices range from 0 to 1 but capture different aspects of specialisation. ND quantifies how many plant species a frugivore interacts with, relative to the total number of fruiting tree species observed at a site, and represents dietary niche breadth (Mello *et al*., 2011). In contrast, Blüthgen’s *d*′ accounts for both interaction frequency and partner availability (the relative availability of each potential interaction partner in the network), and represents complementary specialisation (Blüthgen *et al*., 2006; Carlo *et al*., 2025). Higher ND values indicate greater dietary generalisation and lower values indicate greater specialisation, whereas for *d’*, higher values indicate greater complementary specialisation and vice versa. To examine whether frugivore dietary specialisation varied with frugivore species richness across sites, we checked Pearson’s correlations (since data were normally distributed) between each specialisation index (ND and *d’*) and frugivore species richness. Additionally, we ran GLMMs to model species ND and *d’* as a function of assemblage-level frugivore species richness, with site as a random effect, to test whether richness-specialisation relationships detected at the site level were robust at the species level while accounting for non-independence of species within sites and variation among sites. We used the beta family with “logit” link, as both specialisation indices are proportion values bounded strictly between 0 and 1. The total number of frugivore species included in the analysis was: 26 for Anamalai, 24 for Andaman, 45 for Bala, 40 for Namdapha, seven for Narcondam, and 48 for Pakke.

### Plant-frugivore networks

We constructed weighted plant-frugivore visitation matrices for each study site. The rows of each matrix represented focal fruiting plant species, columns represented avian frugivore species, and the cell entries represented the visitation rate (number of individuals per hour) of each avian frugivore species on each plant species. Most network metrics are sensitive to sampling effort (Blüthgen & Staab, 2024), and the total number of visits we documented across sites varied between 696 and 14000 (Table 1). To overcome the potential influence of variable effort, we used a rarefaction approach as has been recommended (Blüthgen & Staab, 2024). We generated 100 rarefied plant-frugivore visitation matrices per site by randomly sampling 696 interactions (equal to the lowest observed number among our sites) from the original visitation matrix, and for each matrix, calculating the four network metrics, i.e., connectance, nestedness (weighted Nestedness based on Overlap and Decreasing Fill (NODF)), modularity (*Q*), and specialisation (*H2*′) using the R-package ‘bipartite’ (Dormann *et al*., 2009). Each of these network metrics captures distinct ecological properties: connectance measures the proportion of realised links in the network, nestedness indicates whether specialists mostly interact with generalist partners and reflects network redundancy and stability against extinctions, modularity quantifies the degree of compartmentalisation of the network (i.e., cohesive subgroups or modules), and *H2*′ is an index of community-level specialisation (Bascompte *et al*., 2003; Olesen *et al*., 2007; Dormann *et al*., 2009). To examine how these network properties varied with frugivore species richness, we calculated the mean of each rarefield network metric per site and tested their relationship with frugivore species richness using Pearson’s correlations (since data were normally distributed). The number of frugivore species included in the analysis was the same as in the previous section.

All the analyses in this study were done in R (ver. 4.4.1) (R Core Team, 2021).

## RESULTS

We recorded a total of 34,593 visits by 138 species of avian frugivores to 133 species of fruiting plants, resulting in 1,112 unique pairwise plant-frugivore interactions across six assemblages in our study (Table 1).

### Frugivore occurrence on fruiting plants

The mean potential scale reduction factor (PSRF) values for β-parameters (representing species responses to predictors) were 1.002 for Andaman and Pakke, and 1.003 for Anamalai, NTR, and Narcondam, suggesting satisfactory MCMC chain convergence of the HMSC models for all the sites in our study. The mean Tjur *R^2^* values (reflecting the proportion of variation explained by the model) values varied among sites, being lower for the most species-rich sites, (NTR: 0.20; species: 28), and highest for the species-poor site, Narcondam (0.37; species: 5). Variance partitioning among the explanatory variables included in the model showed that the mean variation revealed that, on average across sites, frugivore occurrence on fruiting plants were primarily explained by seed size (mean: 42%), followed by fruit width (23.4%), ripe fruit crop size (19.7%), and pulp lipid content (14.9%) (Fig. 2). This implies morphological traits (fruits and seed traits combined) being prevalent in influencing frugivore occurrence on fruiting plants across south and south-east Asian forests.

**Figure 2.**
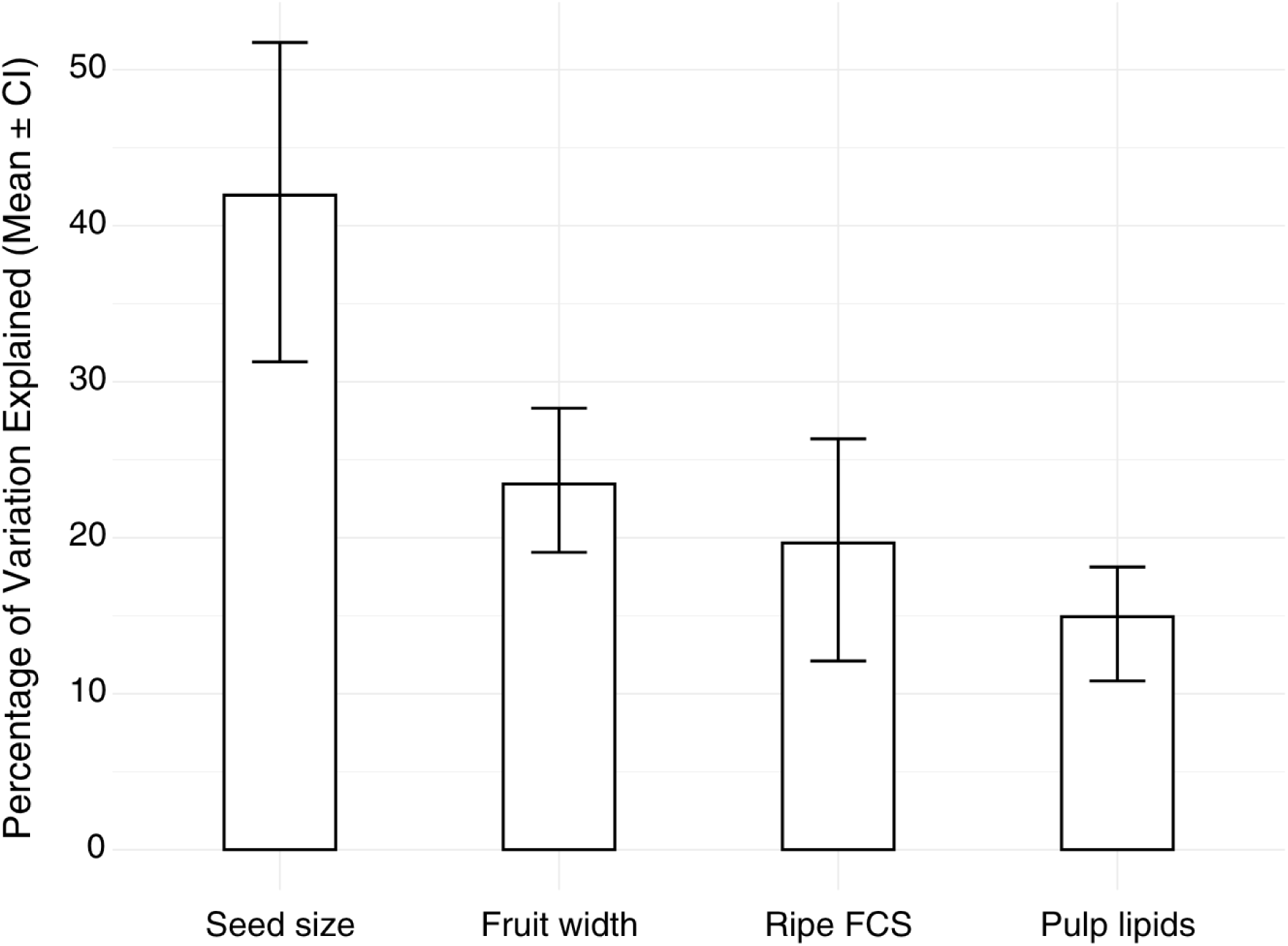
Percentage of variation (Mean ± bootstrapped 95 % CI with 1000 iterations) explained by seed size, fruit width, ripe FCS (fruit crop size, i.e., number of ripe fruits present on a focal plant), and pulp lipids to frugivore occurrence on fruiting plants.

We found statistically significant phylogenetic signals in frugivore responses to predictors for the two species-rich sites (Namdapha and Pakke) but not for species-poor sites (Table S1). However, we did not find a consistent relationship between traits and predictors across sites (Fig. S2). Beak width did not show any association with predictors for the island sites, but was positively associated with fruits with medium lipid content in Anamalai and with fruit width in Namdapha (Fig. S2). Hand-wing index was positively associated with medium seed size, and negatively with medium lipid content in Andaman (Fig. S2). We did not find any statistically significant relationship between traits and predictors for Narcondam and Pakke (Fig. S2).

### Frugivore dietary specialisation

The two measures of dietary specialisation, Normalised Degree (ND) and Blüthgen’s *d*′, yielded different results. The mean ND of avian frugivores ranged between 0.16 (Pakke) and 0.33 (Narcondam) across study sites. Mean ND decreased significantly with increasing frugivore species richness (Pearson’s *r* = –0.87, *p* = 0.024; df = 5; Fig. 3a), indicating a narrower dietary breadth among avian frugivores in species-rich communities. The GLMM results showed a similar trend, with the coefficient being negative and statistically significant at *p* = 0.05 (Fig. S3a). However, the mean *d*′ (range: 0.10 to 0.30 in Anamalai and Pakke, respectively) showed no significant correlation with frugivore species richness (Pearson’s *r* = – 0.03, *p* = 0.949; df = 5; Fig. 3b, S3b), suggesting little evidence for complementary specialisation among frugivores in more species-rich assemblages.

**Figure 3.**
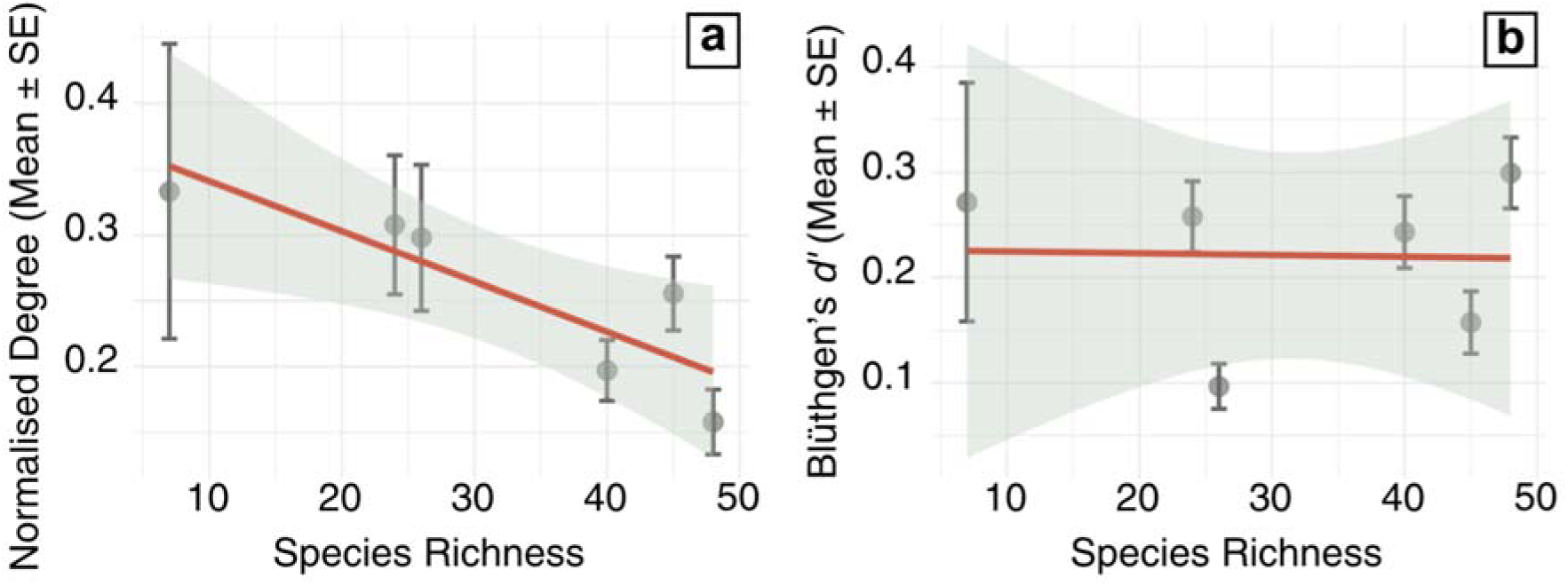
The relationship between Normalised Degree (a) and Blüthgen’s *d*′ (b) with site-level frugivore species richness. The x-axis represents frugivore species richness in our study sites, and the y-axis represents the mean (± standard error) of network metrics. The fitted line in each panel is the regression line, with the shaded area the 95% confidence interval.

### Plant-frugivore networks

Rarefied network metrics varied across sites, with mean values of connectance ranging from 0.12 to 0.33, nestedness from 14.62 to 44.92, modularity from 0.13 to 0.52, and network-level specialisation (*H ^’^*) from 0.23 to 0.52. Mean network connectance and nestedness decreased significantly (*p* < 0.05) with increasing frugivore species richness in the community (Pearson’s *r* = –0.90 and –0.98, respectively; df = 5; Fig. 4a,b). In contrast, network modularity increased significantly with frugivore species richness (Pearson’s *r* = 0.84, *p* < 0.05; Fig. 4c). Although network specialisation was positively correlated with frugivore species richness (Pearson’s *r* = 0.77, *p* = 0.07; Fig. 4d), the relationship was not significant at the α = 0.05 level.

**Figure 4.**
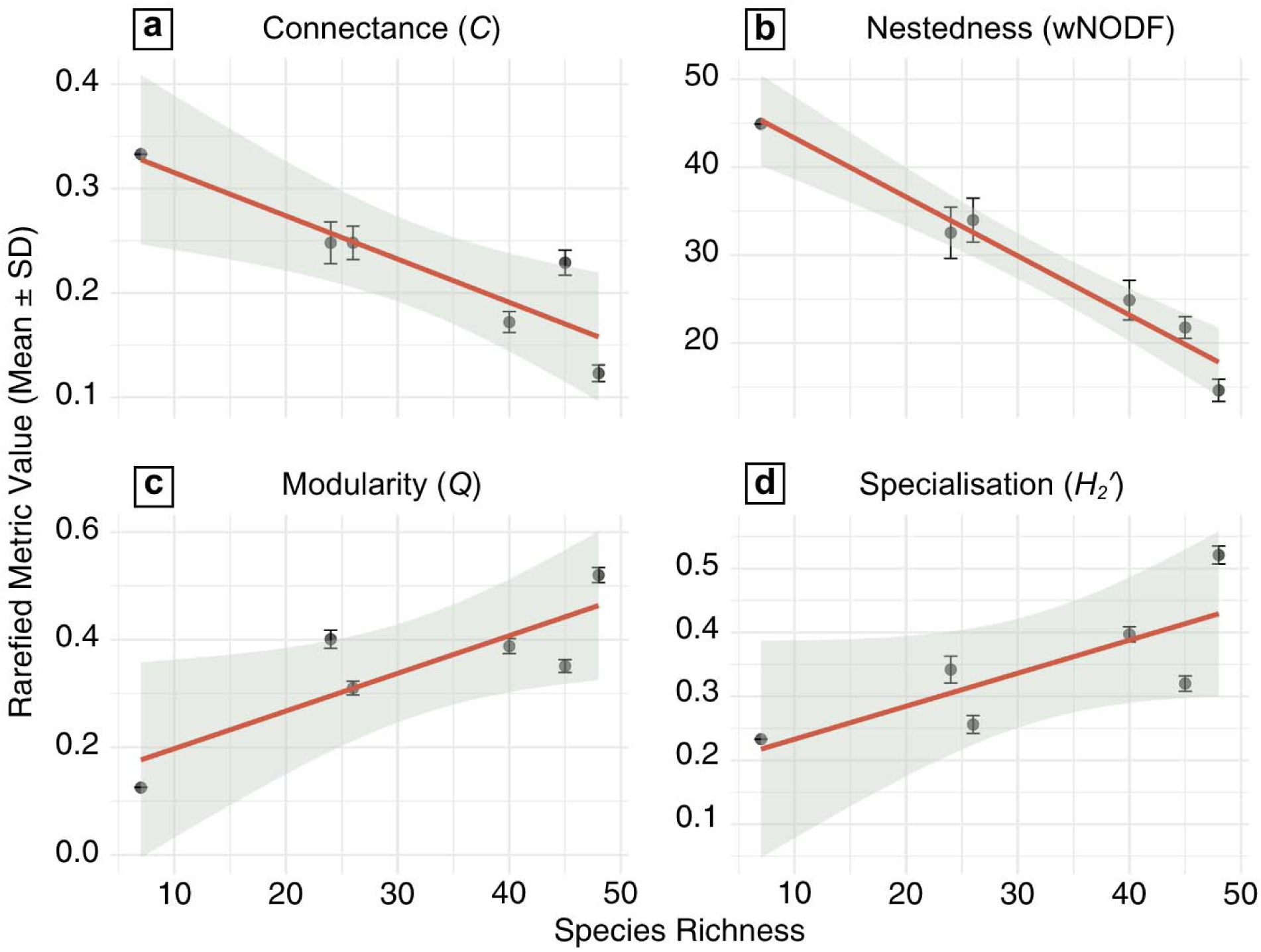
The relationship between the network metrics (a – Connectance, b – Nestedness, c – Modularity, and d – Specialisation) and site-level frugivore species richness. The x-axis represents frugivore species richness in our study sites, and the y-axis represents the mean (± standard deviation) of network metrics calculated using 100 rarefied networks. The fitted line in each panel is the regression line, with the shaded area the 95% confidence interval.

## DISCUSSION

Using primary data from six wet tropical sites in south and south-east Asia, we show that higher frugivore species richness results in increased dietary specialisation, likely leading to cascading effects on network structure: reduced connectance and nestedness, and increased modularity.

Among fruit traits, morphological matching (fruit and seed size) exerted the strongest influence on frugivore occurrence on fruiting plants, outweighing fruit crop size and nutritional traits.

Incidentally, the proportion of explained variation in the occurrence of frugivores on fruiting plants (reflected by Tjur *R^2^* values) declined with increasing frugivore species richness, suggesting that stronger interspecific competition and dietary specialisation in species-rich assemblages reduced the influence of fruit traits. Collectively, our results provide rare empirical evidence that species richness (possibly indicating interspecific competition) likely shapes dietary specialisation and reorganises plant-frugivore networks.

### Species richness, dietary specialisation and network structure

Community assembly processes, like biotic interactions, have been demonstrated to play a role in influencing species richness-niche breadth relationships (Granot & Belmaker, 2020). Consistent with the predictions of the theory of limiting similarity (MacArthur & Levins, 1967), we found that increasing species richness was associated with a reduction in the normalised degree, suggesting narrower dietary niche breadth, i.e., dietary niche specialisation among frugivores. Interestingly, this is in contrast to findings from Neotropics that found greater dietary overlap with increasing niche packing (Dehling *et al*., 2022). While similar examples in the frugivory context are rare, in the pollination context, a study found that with increasing species richness of bees, inter-specific dietary overlap reduced among co-occurring bee species, suggesting niche shifts and dietary specialisation (Fründ *et al*., 2013).

Complementary specialisation (*d*’), a measure of mutual exclusiveness in interactions, did not show any trend with increasing species richness. While very few studies have explicitly examined the relationship between richness, specialisation and network organisation, a recent flower-pollinator study from the Mediterranean region of Spain found that, across a richness gradient, increasing flower and pollinator diversity led to both reduced dietary overlap among pollinators and increased complementary specialisation (*d’*) of pollinators (Gómez-Martínez et al., 2022). In the context of pollination, complementary specialisation may reduce pollen loss and enhance pollen fidelity, thereby benefiting flowers. However, in the context of frugivory, wherein the interactions are more diffused, complementary specialisation may not be expected. Different frugivore species provide complementary seed-dispersal services to plants (Jordano *et al*., 2007), and plants may benefit from being visited by a diverse array of frugivore species (Schupp *et al*., 2010). Existing evidence suggests the absence of phylogenetic signal in fruit colour, indicating that distantly-related plants may have similar fruit colour to attract similar frugivores for seed dispersal, suggesting lowered prevalence of complementary specialisation in plant-frugivore systems (Valenta *et al*., 2018).

While the pollinator richness was explicitly linked with floral richness (Gómez-Martínez *et al*., 2022), we did not find a statistically significant relationship between the number of fruiting plant species observed (i.e., tree species bearing ripe fruits during sampling) and frugivore richness across the same sites (please see Naniwadekar *et al*., 2025 for details). This could be a likely consequence of different mechanisms influencing plant and frugivore communities. Frugivore communities are being assembled due to dispersal limitation over historical time scales, as seen in other studies (Cibois *et al*., 2017). While previous studies have suggested co-diversification between certain fleshy-fruited plant lineages and frugivorous birds such as hornbills (Viseshakul *et al*., 2011), our understanding of plant community assembly in the region remains poor. Limited evidence suggests that regions like the Western Ghats harbour both old and young plant lineages (Gopal *et al*., 2023). However, plant community assembly from an evolutionary lens is yet to be studied across regions.

Previous studies have found contrasting patterns of modularity and nestedness with increasing network complexity. For example, Sebastián González *et al*., (2015) reported that both nestedness and modularity increased with network size, i.e., the total number of interacting plant and frugivore species in the network. In contrast, Almeida & Mikich, (2018) found that modularity decreased with increasing frugivore richness, whereas nestedness showed no significant association with it. Unlike other studies that were conducted across diverse environmental conditions, our study focused on phylogenetically nested plant-frugivore communities experiencing similar climatic conditions. The plant-frugivore communities became more modular and less connected and nested as frugivore richness increased. Thus, network properties can be an outcome of the organisation of interactions as a consequence of biotic factors like inter-specific competition. Our expectation was linked to reduced dietary niche width in species-rich sites, as has been discussed earlier, likely resulting in more compartmentalised communities.

Sampling effort is known to influence network metrics (Blüthgen & Staab, 2024). However, our findings were not influenced by sampling efforts, because the pattern persisted after controlling for sampling effort. Modularity is expected to buffer the community from extinction cascades but reduce functional redundancy (Olesen *et al*., 2007). However, existing evidence from the region suggests that frugivores may exhibit dietary plasticity as dietary shifts have been documented in the absence of competitors (Naniwadekar *et al*., 2021). This suggests that modularity may not affect functional redundancy in these communities. In summary, our findings suggest that increasing frugivore richness drives dietary specialisation and reconfigures network structure, favouring compartmentalisation, without necessarily increasing the exclusivity of interactions.

### Predictors of frugivore visitation

The study also identified the relative influence of different fruit traits on frugivore occurrence across sites with varying species richness (Fig. S1a–e). Given that fruits are patchily distributed in tropical forests and frugivores track these resources, frugivores face contrasting pressures exerted by various fruit traits and availability on one hand, and from interspecific competition on the other. From an optimal foraging perspective, they must balance these contrasting pressures. This study addresses the knowledge gap of the relative importance of different attributes (morphological traits, fruit crop size and fruit nutrients) in tropical assemblages and shows that morphological traits, particularly seed size and fruit width, consistently explained the largest share of variation in frugivore occurrence on fruiting plants across five sites, suggesting a dominant role of morphological trait matching in plant-frugivore community organisation. Fruit and seed size can affect the amount of time the frugivores invest in fruit handling (Palacio *et al*., 2017). For instance, large-bodied birds will be able to swallow small to large fruits with relative ease and handle more fruits, unlike small-bodied frugivores, which may have longer handling times. Consequently, morphological traits can influence frugivore visitation, and over evolutionary time scales, frugivore assemblages can reciprocally influence fruit and seed traits through selection pressures imposed by their feeding preferences and gape limitations (Valenta & Nevo, 2020). However, there appears to be a reducing influence of seed size (but not other traits) in explaining the occurrence of frugivores on fruiting trees. Moreover, the explained variation in the occurrence of frugivores was the least for the species-rich sites and highest for the species-poor sites. This can be an outcome of the influence of biotic interactions (inter-specific competition) in influencing frugivore visitations on fruiting trees. According to the dispersal syndrome hypothesis, seed dispersers exert selection pressures on fruit traits promoting morphological convergence towards potentially improved seed dispersal outcomes (Galetti *et al*., 2013). The patterns of increased dietary and network specialisation observed in this study underscore the potential importance of biotic interactions in shaping such evolutionary outcomes. Future studies should test how variation in frugivore assemblage composition and trait diversity translates into selection on fruit and seed traits across tropical forest communities.

The influence of fruit nutrients on influencing frugivore visitations on fruiting plants may differ. For instance, insectivorous and omnivorous birds have been demonstrated to strongly track lipid content in fruits, vis-à-vis frugivorous birds (Pizo *et al*., 2021). Moreover, frugivores may consume a combination of lipid-rich and poor diets to balance the uptake of different macronutrients (Blendinger *et al*., 2022). This likely resulted in our observed pattern of relatively lower influence of lipid content in fruits on the occurrence of frugivores on fruiting plants.

Interestingly, we did not find consistent associations between frugivore traits (beak width, hand-wing index, degree of frugivory) and fruit or seed traits, fruit crop size, or pulp nutrient across the six sites (Fig. S2). This multi-site comparison, spanning a gradient in frugivore species richness, suggests an absence of generalisable, cross-site trait-predictor relationships governing frugivore occurrence on fruiting plants. This pattern likely highlights the strong context dependence of trait-resource relationships in frugivore assemblages. Frugivores in tropical forests are known to exhibit substantial dietary plasticity, adjusting diets based on the presence or absence of sympatric frugivores (Naniwadekar *et al*., 2021). Such plasticity has been documented in several tropical bird assemblages, where species shift diets when competitors are absent but exhibit altered diets in species-rich communities. This behavioural plasticity likely weakens trait-predictor associations across sites. Interestingly, we detected significant phylogenetic signal in species responses to fruit traits at the two most species-rich sites. Species-rich assemblages in our study are characterised by greater representation within lineages rather than by the presence of many distantly-related lineages. Closely-related species often have similar morphological traits and may therefore respond similarly to fruit traits, leading to phylogenetic signal in resource tracking even after accounting for measured functional traits. However, this needs further exploration as studies from the Eastern Himalaya have shown that sympatric hornbill species differ in their reliance on figs versus non-fig fruits, potentially facilitating coexistence through dietary differentiation. If this is a pervasive pattern across lineages, then phylogenetic signal may not be expected. Absence of similar information from other lineages precludes such examination, an aspect that needs to be explored in future studies.

### Limitations and future directions

The observed patterns of dietary niche specialisation, decreased connectance, nestedness, and increased modularity with increasing frugivore species richness are based on data from only six sites. However, comparable datasets that deploy consistent methods to quantify plant-frugivore interactions across species richness gradients, while maintaining broadly similar environmental regimes and nested phylogenetic composition of frugivore assemblages, are extremely rare. We therefore exercised caution by employing rarefaction (for network metrics) and multiple analytical frameworks (for normalised degree and dietary specialisation) to validate our conclusions. While direct evidence of interspecific competition is difficult to obtain in field settings, several studies suggest that increasing species richness can intensify interspecific competition and promote greater niche specialisation (Freeman *et al*., 2024). Finally, while this study examined the influence of species richness on the dietary niche of frugivores, future studies should examine how richness influences resource partitioning in other important niche dimensions, such as space and time.

## CONCLUSIONS

We provide evidence for increased dietary specialisation in communities harbouring higher species richness, with consequences for understanding the organisation of communities characterised by mutualistic interactions. This suggests increasing species richness results in increased niche specialisation, with communities becoming more modular and less nested and connected. Unlike in the pollination context, in frugivory, niche specialisation is not necessarily accompanied by complementary specialisation, as frugivores may provide complementary services to plants. We also highlight the relatively important role of fruit and seed morphology in influencing interactions between plants and frugivores. Our results are among the few field-based studies that control for environmental and phylogenetic effects while examining the richness effects on the dietary niches of frugivores and community organisation.

## ETHICS APPROVAL STATEMENT

This was an observational study that did not require handling of animals. The authors got permissions from the respective Forest Departments before conducting the study.

## ACKNOWLEDGEMENTS

We thank the Forest Departments of Arunachal Pradesh, Andaman and Nicobar Islands, Tamil Nadu, and the Thailand Department of National Park, Wildlife and Plant Conservation. We thank George Gale, Aparajita Datta, Divya Mudappa, T. R. Shankar Raman, Mousumi Ghosh, M Ananda Kumar, Japang Pansa, Hasan Ali, Late Josna Malik, Devathi Parashuram, Daphawan Khamcha, Sitthichai Jinamoy, Siriwan Nakkuntod, Sunate Karapan, Sukanya Chaisuriyanun, Vignesh Chandran, and Narongsak Pongdee for support and critical discussions. We thank our field staff across the different field sites: Pakke: Khem Thapa, Late Tali Nabam, Late Kumar Thapa; Andaman: Michael Kujur, Sujith Bengra; Namdapha: Dhan Bahadur Limbu, Wanggao Wangsa, Laiphong Wangnao; Anamalai: Krishnakumar, Sathiyaraj; and Bala: Isamaea Ma, Iswan Jowae, Masauphee Hub. We thank Siddhant Mehendale, Prabhav Benara, and Mandar Tijare for assisting us in data collection in Anamalai.

