## Supplementary material for "Frugivore species richness influences dietary specialisation and network properties in Asian wet tropical forests": Fig. S; Table S

**Table S1.** The posterior distribution of the phylogenetic signal parameter *ρ* (Mean and 95% credible intervals) for the five sites used for the Hierarchical Modelling of Species Communities (HMSC) analysis. The value of *ρ* ranges between 0 (no phylogenetic signal) and 1 (very strong phylogenetic signal).

| **Site** | **Mean *ρ* (95% CI)** |
| --- | --- |
| Narcondam | 0.36 (0.00–0.98) |
| Andaman | 0.08 (0.00–0.61) |
| Anamalai | 0.38 (0.00–0.91) |
| Pakke | 0.88 (0.42–1.00) |
| Namdapha | 0.79 (0.24–0.99) |

**Table S2.** Information on fruit and seed size, and pulp nutrient content of focal plant species across the study sites. Fruit width was measured in the field using a Mitutoyo digital vernier calliper. Seed width was also measured for most species; however, for a few species, seed size was subsequently categorised using expert knowledge and existing databases. Pulp lipid content was estimated for 18 of the 28 fruiting plant species at the Anamalai Rainforest Research Station (hereafter Anamalai), owing to better fruit-processing facilities and access to a laboratory. Lipid information for the remaining species was collated from published literature at the finest taxonomic resolution possible (i.e., species, genus, or family).

| **Sl. no.** | **Plant species** | **Site(s) present in** | **Seed size (categories)** | **Fruit width (mm)** | **Pulp lipids (%)** | **Pulp lipids (categories)** | **Taxonomic levels of pulp lipid estimation** | **Reference(s) for pulp lipid information** |
| --- | --- | --- | --- | --- | --- | --- | --- | --- |
| 1 | *Actinodaphne malabarica* | Anamalai | Medium | 9.14 | 7.12 | Low | Species | Mandal *et al*., 2026 |
| 2 | *Actinodaphne obovata* | Pakke | Medium | NA | NA | NA | NA | NA |
| 3 | *Aglaia* sp. Bala | Bala | Large | NA | NA | NA | NA | NA |
| 4 | *Aglaia* sp. Pakke | Pakke | Large | 19.64 | 25.99 | Medium | Genus | Jordano, 1995; Savini, 2007 |
| 5 | *Aglaia spectabilis* | Pakke | Medium | 16.46 | 25.99 | Medium | Genus | Jordano, 1995; Savini, 2007 |
| 6 | Aidia densiflora | Narcondam | Small | 7.68 | 4.64 | Low | Family | Jordano, 1995 |
| 7 | *Alseodaphne* sp. Namdapha | Namdapha | Large | 23.39 | 24.61 | Medium | Family | Corlett, 1996; Jordano, 1995; Savini, 2007; Mandal *et al*., 2026; Pizo *et al*., 2021 |
| 8 | *Anamirta cocculus* | Narcondam | Medium | 13.11 | 3.45 | Low | Family | Jordano, 1995 |
| 9 | *Antidesma montanum* | Pakke | Small | 5 | 4.17 | Low | Genus | Corlett, 1996; Wilson & Downs, 2012 |
| 10 | *Antidesma velutinosum* | Bala | Small | NA | NA | NA | NA | NA |
| 11 | *Aphanamixis polystachya* | Andaman | Large | 16.03 | 31.67 | Medium | Family | Beehler & Dumbacher, 1996; Foster & McDiarmid, 1983; Savini 2007; Galetti *et al*., 2011; Jordano, 1995; Kalenga Saka & Msonthi, 1994; Wilson & Downs, 2012 |
| 12 | *Aporosa octandra* | Andaman | Medium | 10.95 | 1.00 | Low | Genus | Corlett, 1996 |
| 13 | *Aralia* sp. Namdapha | Namdapha | Small | 4.32 | 2.00 | Low | Genus | Corlett, 1996 |
| 14 | *Beilschmiedia assamica* | Namdapha, Pakke | Large | 23.14 | 12.95 | Medium | Genus | Jordano, 1995; Savini, 2007 |
| 15 | *Beilschmiedia* sp. Namdapha | Namdapha | Large | 16.99 | 12.95 | Medium | Genus | Jordano, 1995; Savini, 2007 |
| 16 | *Beilschmiedia* sp. Pakke 1 | Pakke | Medium | 14.76 | 12.95 | Medium | Genus | Jordano, 1995; Savini, 2007 |
| 17 | *Beilschmiedia* sp. Pakke 2 | Pakke | Large | 20.3 | 12.95 | Medium | Genus | Jordano, 1995; Savini, 2007 |
| 18 | *Bischofia javanica* | Anamalai, Namdapha, Pakke | Small | 10.44/9.29/9.92 | 1.35 | Low | Genus | Corlett, 1996; Suwardi *et al*., 2022; Mandal *et al*., 2026 |
| 19 | *Bridelia glauca* | Pakke | Small | 8.82 | 1.43 | Low | Genus | Corlett, 1996; Jordano, 1995; Savini, 2007; Wilson & Downs, 2012 |
| 20 | *Bridelia retusa* | Anamalai, Andaman | Medium/Small | 10.01/5.64 | 1.43 | Low | Genus | Corlett, 1996; Jordano, 1995; Savini, 2007; Wilson & Downs, 2012 |
| 21 | *Bridelia squamosa* | Anamalai | Small | 6.72 | 1.43 | Low | Genus | Corlett, 1996; Jordano, 1995; Savini, 2007; Wilson & Downs, 2012 |
| 22 | *Bridelia tomentosa* | Bala | Small | NA | NA | NA | NA | NA |
| 23 | *Buchnania* sp. Andaman | Andaman | NA | NA | NA | NA | NA | NA |
| 24 | *Callicarpa arborea* | Pakke | Small | 3 | 1.50 | Low | Genus | Corlett, 1996 |
| 25 | *Campnosperma auriculatum* | Bala | Small | NA | NA | NA | NA | NA |
| 26 | *Canarium euphyllum* | Andaman, Narcondam | Medium | NA | NA | NA | NA | NA |
| 27 | *Canarium strictum* | Anamalai, Namdapha | Large | 26.25/23.22 | 20.23 | Medium | Genus | Jordano, 1995; Savini, 2007; Sweeney *et al*., 2017; Mandal *et al*., 2026 |
| 28 | *Carallia brachiata* | Pakke | Medium | 9.38 | 7.00 | Low | Species | Corlett, 1996 |
| 29 | *Caryota mitis* | Narcondam | Large | 15.61 | 1.50 | Low | Genus | Perumpuli *et al*., 2022 |
| 30 | *Chionanthus macrophyllus* | Namdapha | Medium | 10.98 | 1.90 | Low | Genus | Jordano, 1995 |
| 31 | *Chionanthus* sp. Namdapha | Namdapha | Large | 28.25 | 1.90 | Low | Genus | Jordano, 1995 |
| 32 | *Chionanthus* sp. Narcondam | Narcondam | Medium | 14 | 1.90 | Low | Genus | Jordano, 1995 |
| 33 | *Chisocheton ceramicus* | Bala | Large | NA | NA | NA | NA | NA |
| 34 | *Chisocheton cumingianus* | Pakke | Large | 23.34 | 60.00 | High | Genus | Beehler & Dumbacher, 1996 |
| 35 | *Cinnamomum bejolghota* | Pakke | Medium | 8.08 | 25.77 | Medium | Genus | Corlett, 1996; Jordano, 1995; Savini, 2007; Pizo *et al*., 2021 |
| 36 | *Cissus repanda* | Anamalai | Medium | 6.27 | 14.37 | Medium | Genus | Jordano, 1995 |
| 37 | *Codiocarpus andamanicus* | Narcondam | NA | NA | NA | NA | NA | NA |
| 38 | *Debregeasia* sp. Andaman | Andaman | Small | 6.55 | 2.39 | Low | Genus | Seal & Chaudhuri, 2014 |
| 39 | *Dendrocnide sinuata* | Pakke | Small | 4.5 | 5.98 | Low | Family | Jordano, 1995; Seal & Chaudhuri, 2014 |
| 40 | *Dendrocnide* sp. Namdapha | Namdapha | Small | 3.03 | 5.98 | Low | Family | Jordano, 1995; Seal & Chaudhuri, 2014 |
| 41 | *Dysoxylum cauliflorum* | Pakke | Medium | 12.67 | 37.18 | High | Genus | Beehler & Dumbacher, 1996; Jordano, 1995; Savini, 2007 |
| 42 | *Dysoxylum gotadhora* | Pakke | Large | 21.36 | 37.18 | High | Genus | Beehler & Dumbacher, 1996; Jordano, 1995; Savini, 2007 |
| 43 | *Dysoxylum* sp. Andaman | Andaman | Large | 22.43 | 37.18 | High | Genus | Beehler & Dumbacher, 1996; Jordano, 1995; Savini, 2007 |
| 44 | *Dysoxylum* sp. Namdapha | Namdapha | Large | 20.5 | 37.18 | High | Genus | Beehler & Dumbacher, 1996; Jordano, 1995; Savini, 2007 |
| 45 | *Ehretia wallichiana* | Pakke | Small | 7 | 2.20 | Low | Genus | Jordano, 1995 |
| 46 | *Elaeocarpus angustifolius* | Pakke | Large | 25.79 | 10.86 | Medium | Genus | Corlett, 1996; Jordano, 1995; Mandal *et al*., 2026 |
| 47 | *Elaeocarpus palembanicus* | Bala | Medium | NA | NA | NA | NA | NA |
| 48 | *Elaeocarpus stipularis* | Bala | Large | NA | NA | NA | NA | NA |
| 49 | *Endocomia macrocoma* | Narcondam | Large | 28.67 | 38.36 | High | Family | Beehler & Dumbacher, 1996; Galetti et al., 2011; Jordano, 1995; Savini, 2007; Mandal *et al*., 2026 |
| 50 | *Endocomia* sp. Andaman | Andaman | Large | 17.93 | 38.36 | High | Family | Beehler & Dumbacher, 1996; Galetti *et al*., 2011; Jordano, 1995; Savini, 2007; Mandal *et al*., 2026 |
| 51 | *Eriobotrya bengalensis* | Bala | Small | NA | NA | NA | NA | NA |
| 52 | *Falconeria insignis* | Pakke | Medium | 7.91 | 35.14 | High | Family | Corlett, 1996; Jordano, 1995; Galetti *et al*., 2011 |
| 53 | *Ficus altissima* | Andaman, Namdapha, Pakke | Small | 17.12/18.29/18.84 | 3.37 | Low | Genus | Corlett, 1996; Jordano, 1995; Savini, 2007; Mandal *et al*., 2026; Onrizal & Auliah, 2019; Wilson & Downs, 2012 |
| 54 | *Ficus amplissima* | Anamalai | Small | 9.53 | 1.02 | Low | Species | Mandal *et al*., 2026 |
| 55 | *Ficus benjamina* | Andaman, Narcondam, Pakke | Small | 16.26/11.61/16.26 | 3.37 | Low | Genus | Corlett, 1996; Jordano, 1995; Savini, 2007; Mandal *et al*., 2026; Onrizal & Auliah, 2019; Wilson & Downs, 2012 |
| 56 | *Ficus costata* | Andaman | Small | 3.47 | 3.37 | Low | Genus | Corlett, 1996; Jordano, 1995; Savini, 2007; Mandal *et al*., 2026; Onrizal & Auliah, 2019; Wilson & Downs, 2012 |
| 57 | *Ficus cucurbitina* | Bala | Small | NA | NA | NA | NA | NA |
| 58 | *Ficus drupacea* | Anamalai, Namdapha, Pakke | Small | 19.41/37.68/31.63 | 2.98 | Low | Species | Mandal *et al*., 2026 |
| 59 | *Ficus dubia* | Bala | Small | NA | NA | NA | NA | NA |
| 60 | *Ficus exasperata* | Anamalai | Small | 19.94 | 1.34 | Low | Species | Mandal *et al*., 2026 |
| 61 | *Ficus geniculata* | Andaman, Namdapha, Pakke | Small | 5.71/9.48/6.94 | 3.37 | Low | Genus | Corlett, 1996; Jordano, 1995; Savini, 2007; Mandal *et al*., 2026; Onrizal & Auliah, 2019; Wilson & Downs, 2012 |
| 62 | *Ficus glaberima* | Narcondam | Small | 13.31 | 3.37 | Low | Genus | Corlett, 1996; Jordano, 1995; Savini, 2007; Mandal *et al*., 2026; Onrizal & Auliah, 2019; Wilson & Downs, 2012 |
| 63 | *Ficus microcarpa* | Anamalai | Small | 8.52 | 0.32 | Low | Species | Mandal *et al*., 2026 |
| 64 | *Ficus nervosa* | Anamalai, Namdapha, Narcondam, Pakke | Small | 18.23/9.83/21.89/10.85 | 0.78 | Low | Species | Mandal *et al*., 2026 |
| 65 | *Ficus obtusifolia* | Andaman, Pakke | Small | 16.06/12.53 | 3.37 | Low | Genus | Corlett, 1996; Jordano, 1995; Savini, 2007; Mandal *et al*., 2026; Onrizal & Auliah, 2019; Wilson & Downs, 2012 |
| 66 | *Ficus rumphii* | Andaman, Narcondam | Small | 10.96/11.06 | 3.37 | Low | Genus | Corlett, 1996; Jordano, 1995; Savini, 2007; Mandal et al., 2026; Onrizal & Auliah, 2019; Wilson & Downs, 2012 |
| 67 | *Ficus scandens* | Namdapha | Small | 6.99 | 3.37 | Low | Genus | Corlett, 1996; Jordano, 1995; Savini, 2007; Mandal *et al*., 2026; Onrizal & Auliah, 2019; Wilson & Downs, 2012 |
| 68 | *Ficus* sp. Andaman | Andaman | Small | NA | NA | NA | NA | NA |
| 69 | *Ficus* sp. Bala | Bala | Small | NA | NA | NA | NA | NA |
| 70 | *Ficus* sp. Namdapha 1 | Namdapha | Small | 8.71 | 3.37 | Low | Genus | Corlett, 1996; Jordano, 1995; Savini, 2007; Mandal *et al*., 2026; Onrizal & Auliah, 2019; Wilson & Downs, 2012 |
| 71 | *Ficus* sp. Namdapha 2 | Namdapha | Small | 14.75 | 3.37 | Low | Genus | Corlett, 1996; Jordano, 1995; Savini, 2007; Mandal *et al*., 2026; Onrizal & Auliah, 2019; Wilson & Downs, 2012 |
| 72 | *Ficus* sp. Pakke | Pakke | Small | 12.12 | 3.37 | Low | Genus | Corlett, 1996; Jordano, 1995; Savini, 2007; Mandal *et al*., 2026; Onrizal & Auliah, 2019; Wilson & Downs, 2012 |
| 73 | *Ficus sumatrana* | Bala | Small | NA | NA | NA | NA | NA |
| 74 | *Ficus sundaica* | Bala | Small | NA | NA | NA | NA | NA |
| 75 | *Ficus superba* | Anamalai, Narcondam | Small | 5.97/6.00 | 3.37 | Low | Genus | Corlett, 1996; Jordano, 1995; Savini, 2007; Mandal *et al*., 2026; Onrizal & Auliah, 2019; Wilson & Downs, 2012 |
| 76 | *Ficus tinctoria* | Anamalai, Andaman, Namdapha | Small | 4.89/9.08/4.7 | 2.18 | Low | Species | Mandal *et al*., 2026 |
| 77 | *Ficus tsjakela* | Anamalai | Small | 4.86 | 1.64 | Low | Species | Mandal *et al*., 2026 |
| 78 | *Ficus virens* | Anamalai, Andaman | Small | 16.82/10.49 | 0.90 | Low | Species | Mandal *et al*., 2026 |
| 79 | *Filicium decipiens* | Anamalai | Medium | 9.96 | 0.60 | Low | Species | Mandal *et al*., 2026 |
| 80 | *Flacourtia montana* | Anamalai | Small | 13.54 | 0.14 | Low | Species | Mandal *et al*., 2026 |
| 81 | *Glochidion obscurum* | Bala | Small | NA | NA | NA | NA | NA |
| 82 | *Glochidion* sp. Bala | Bala | Small | NA | NA | NA | NA | NA |
| 83 | *Gymnacranthera contracta* | Bala | Large | NA | NA | NA | NA | NA |
| 84 | *Gynotroches axillaris* | Bala | Small | NA | NA | NA | NA | NA |
| 85 | *Heteropanax fragrans* | Andaman, Namdapha, Pakke | Small | 9.97/10.78/7.62 | 19.48 | Medium | Family | Corlett, 1996; Jordano, 1995; Pizo *et al*., 2021 |
| 86 | *Heynea trijuga* | Anamalai | Medium | 12.7 | 41.84 | High | Genus | Foster & McDiarmid, 1983; Jordano, 1995; Kalenga Saka & Msonthi, 1994; Wilson & Downs, 2012 |
| 87 | *Horsfieldia kingii* | Namdapha, Pakke | Large | 22.97/20.73 | 41.61 | High | Genus | Savini, 2007 |
| 88 | *Hovenia acerba* | Namdapha | Small | 4.75 | 0.46 | Low | Genus | Maieves *et al*., 2015 |
| 89 | *Ilex umbellulata* | Andamans | Small | 4.93 | 2.05 | Low | Genus | Corlett, 1996; Jordano, 1995; Seal & Chaudhuri, 2014 |
| 90 | *Knema attenuata* | Anamalai | Large | 20.27 | 18.20 | Medium | Genus | Savini, 2007 |
| 91 | *Knema erratica* | Pakke | Medium | 15.16 | 18.20 | Medium | Genus | Savini, 2007 |
| 92 | *Knema furfuracea* | Bala | Medium | NA | NA | NA | NA | NA |
| 93 | *Knema* sp. Andaman | Andaman | Medium | 16.6 | 18.20 | Medium | Genus | Savini, 2007 |
| 94 | *Lannea coromandelica* | Andaman | Medium | 6.92 | 1.87 | Low | Genus | Muhammad *et al*., 2018 |
| 95 | *Leea indica* | Pakke | Small | 7 | 0.95 | Low | Genus | Onrizal & Auliah, 2019 |
| 96 | *Litsea coriacea* | Anamalai | Small | 7.79 | 14.42 | Medium | Species | Mandal *et al*., 2026 |
| 97 | *Litsea* sp. Bala | Bala | Small | NA | NA | NA | NA | NA |
| 98 | *Litsea* sp. Namdapha | Namdapha | Medium | 15.57 | 20.08 | Medium | Genus | Corlett, 1996; Jordano, 1995; Savini, 2007; Mandal *et al*., 2026 |
| 99 | *Litsea* sp. Pakke 1 | Pakke | Medium | NA | NA | NA | NA | NA |
| 100 | *Litsea* sp. Pakke 2 | Pakke | Medium | NA | NA | NA | NA | NA |
| 101 | *Livistona jenkinsiana* | Pakke | Large | 27.75 | 30.61 | Medium | Genus | Savini, 2007 |
| 102 | *Macaranga indica* | Namdapha | Small | 4.09 | 29.00 | Medium | Genus | Corlett, 1996 |
| 103 | *Macaranga peltata* | Anamalai | Small | 5.41 | 29.00 | Medium | Genus | Corlett, 1996 |
| 104 | *Micromelum integerrimum* | Pakke | Medium | 8.75 | 6.51 | Low | Family | Corlett, 1996; Jordano, 1995; Mandal *et al*., 2026 |
| 105 | *Myristica dactyloides* | Anamalai | Large | 29.18 | 44.80 | High | Genus | Jordano, 1995, Beehler, 1996 |
| 106 | *Myristica* sp. Andaman | Andaman | Large | 22.66 | 44.80 | High | Genus | Jordano, 1995, Beehler, 1996 |
| 107 | *Olea dioica* | Anamalai, Pakke | Medium | 9.16/11.1 | 11.76 | Medium | Genus | Jordano, 1995; Mandal *et al*., 2026 |
| 108 | *Persea macrantha* | Anamalai, Namdapha, Pakke | Medium/Large | 15.94/20.24 | 35.03 | High | Genus | Corlett, 1996; Jordano, 1995; Mandal *et al*., 2026; Pizo *et al*., 2021 |
| 109 | *Phoebe* sp. Namdapha | Namdapha | Medium | 26.3 | 26.83 | Medium | Genus | Jordano, 1995; Savini, 2007 |
| 110 | *Phoebe* sp. Pakke 1 | Pakke | Medium | 12.32 | 26.83 | Medium | Genus | Jordano, 1995; Savini, 2007 |
| 111 | *Phoebe* sp. Pakke 2 | Pakke | NA | NA | NA | NA | NA | NA |
| 112 | *Picrasma javanica* | Pakke | Medium | 10.62 | 42.00 | High | Genus | Corlett, 1996 |
| 113 | *Pinanga andamanensis* | Andaman | Medium | 9 | 1.18 | Low | Genus | Riley *et al*., 2013; Silva *et al*., 2015; Sweeney *et al*., 2017 |
| 114 | *Polyalthia fragrans* | Anamalai | Medium | 15.44 | 6.69 | Low | Genus | Jordano, 1995; Savini, 2007; Onrizal & Auliah, 2019 |
| 115 | *Polyalthia simiarum* | Pakke | Medium | 20.94 | 6.69 | Low | Genus | Jordano, 1995; Savini, 2007; Onrizal & Auliah, 2019 |
| 116 | *Prunus ceylanica* | Namdapha, Pakke | Large | 33.15/21.92 | 2.00 | Low | Genus | Jordano, 1995 |
| 117 | *Prunus* sp. Andaman | Andaman | Large | 20.32 | 2.00 | Low | Genus | Jordano, 1995 |
| 118 | *Sapium baccatum* | Narcondam | Small | 12 | 51.47 | High | Genus | Jordano, 1995; Corlett, 1996 |
| 119 | *Sarcotheca laxa* | Bala | Small | NA | NA | NA | NA | NA |
| 120 | *Saurauia* sp. Bala | Bala | Small | NA | NA | NA | NA | NA |
| 121 | *Schefflera* sp. Namdapha | Namdapha | Small | 5.27 | 18.45 | Medium | Genus | Corlett, 1996; Pizo *et al*., 2021 |
| 122 | *Sloanea sterculiaceae* | Pakke | Medium | 9.96 | 2.75 | Low | Genus | Galetti, 2011, Pizo, 2021 |
| 123 | *Sterculia rubiginosa* | Andaman | Medium | 10.63 | 31.00 | Medium | Genus | Corlett, 1996 |
| 124 | *Sterculia villosa* | Pakke | Medium | 6.95 | 31.00 | Medium | Genus | Corlett, 1996 |
| 125 | *Syzygium lanceolatum* | Anamalai | Medium | 9.91 | 2.22 | Low | Genus | Corlett, 1996; Jordano, 1995; Kalenga Saka & Msonthi, 1994; Savini, 2007; Wilson & Downs, 2012; Suwardi *et al*., 2022 |
| 126 | *Syzygium* sp. Pakke | Pakke | Medium | 8.73 | 2.22 | Low | Genus | Corlett, 1996; Jordano, 1995; Kalenga Saka & Msonthi, 1994; Savini, 2007; Wilson & Downs, 2012; Suwardi *et al*., 2022 |
| 127 | *Tetradium glabrifolium* | Pakke | Small | 4 | 6.51 | Low | Family | Corlett, 1996; Jordano, 1995; Mandal *et al*., 2026 |
| 128 | *Trema orientalis* | Namdapha | Small | 2.5 | 40.55 | High | Species | Jordano, 1995; Wilson & Downs, 2012 |
| 129 | Unid. sp. Andaman | Andaman | Medium | 17.61 | NA | NA | NA | NA |
| 130 | *Urophyllum* sp. Bala | Bala | Small | NA | NA | NA | NA | NA |
| 131 | *Villebrunea integrifolia* | Anamalai | Small | 6.38 | NA | NA | NA | NA |
| 132 | *Vitex glabrata* | Pakke | Small | 13 | 0.67 | Low | Genus | Jordano, 1995; Kalenga Saka & Msonthi, 1994; Galetti *et al*., 2011 |
| 133 | *Zanthoxylum rhetsa* | Anamalai, Pakke | Medium | 7.59/7.72 | 1.20 | Low | Species | Mandal *et al*., 2026 |

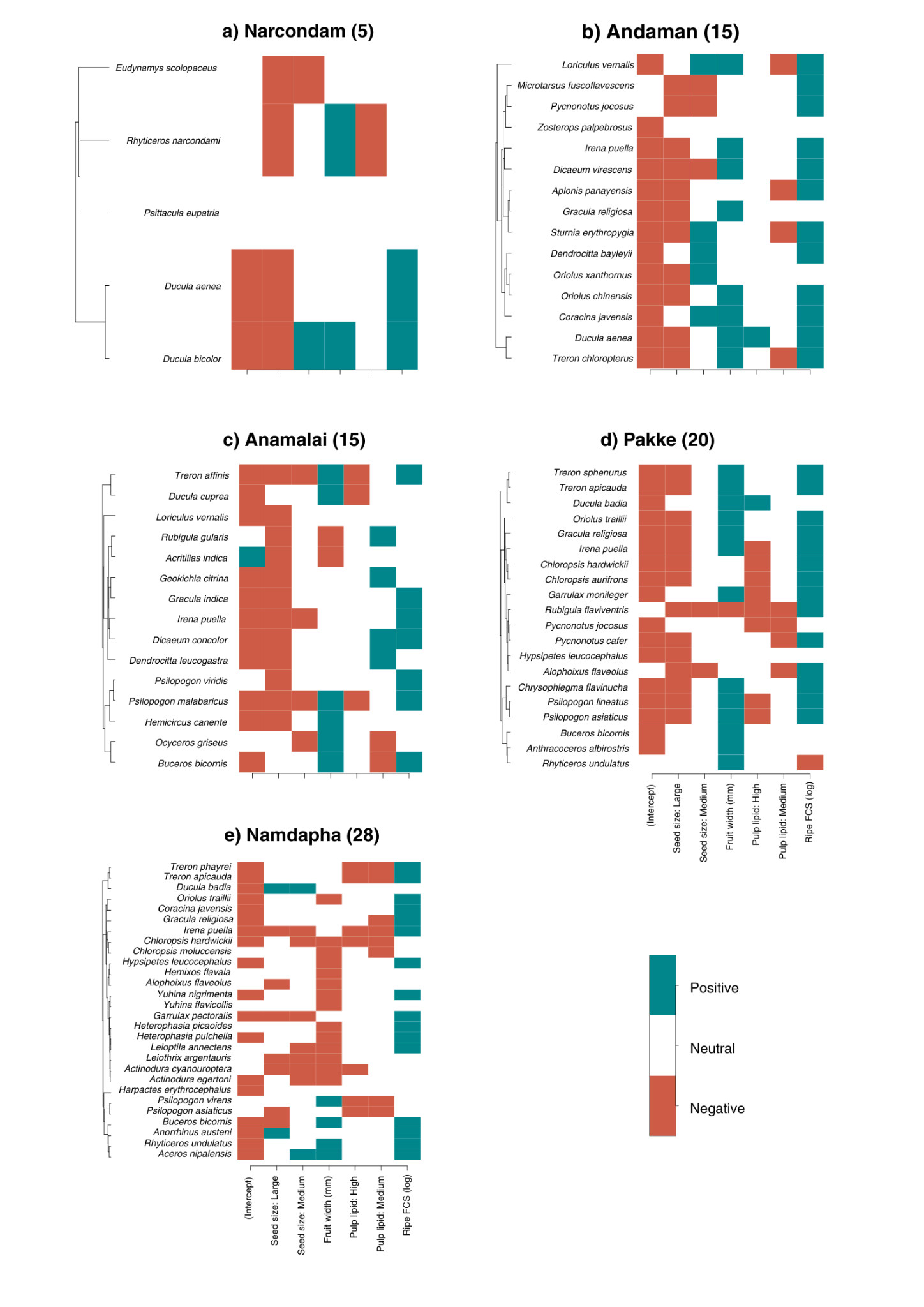

**Fig. S1.** Heatmaps showing species-specific responses to the fixed effects used in the HMSC analysis, namely, seed size (small, medium, and large), fruit width (mm), pulp lipid (low, medium, and high), and log-transformed ripe fruit crop size (FCS) for our five study sites, and how the responses were influenced by their phylogeny. Each panel represent one study site, and the numbers in parentheses correspond to the number of frugivore species used in the analysis for that particular study site. The orange colour represents a positive relationship, turquoise represents a negative relationship, and white represents no relationship at a 95% posterior support level.

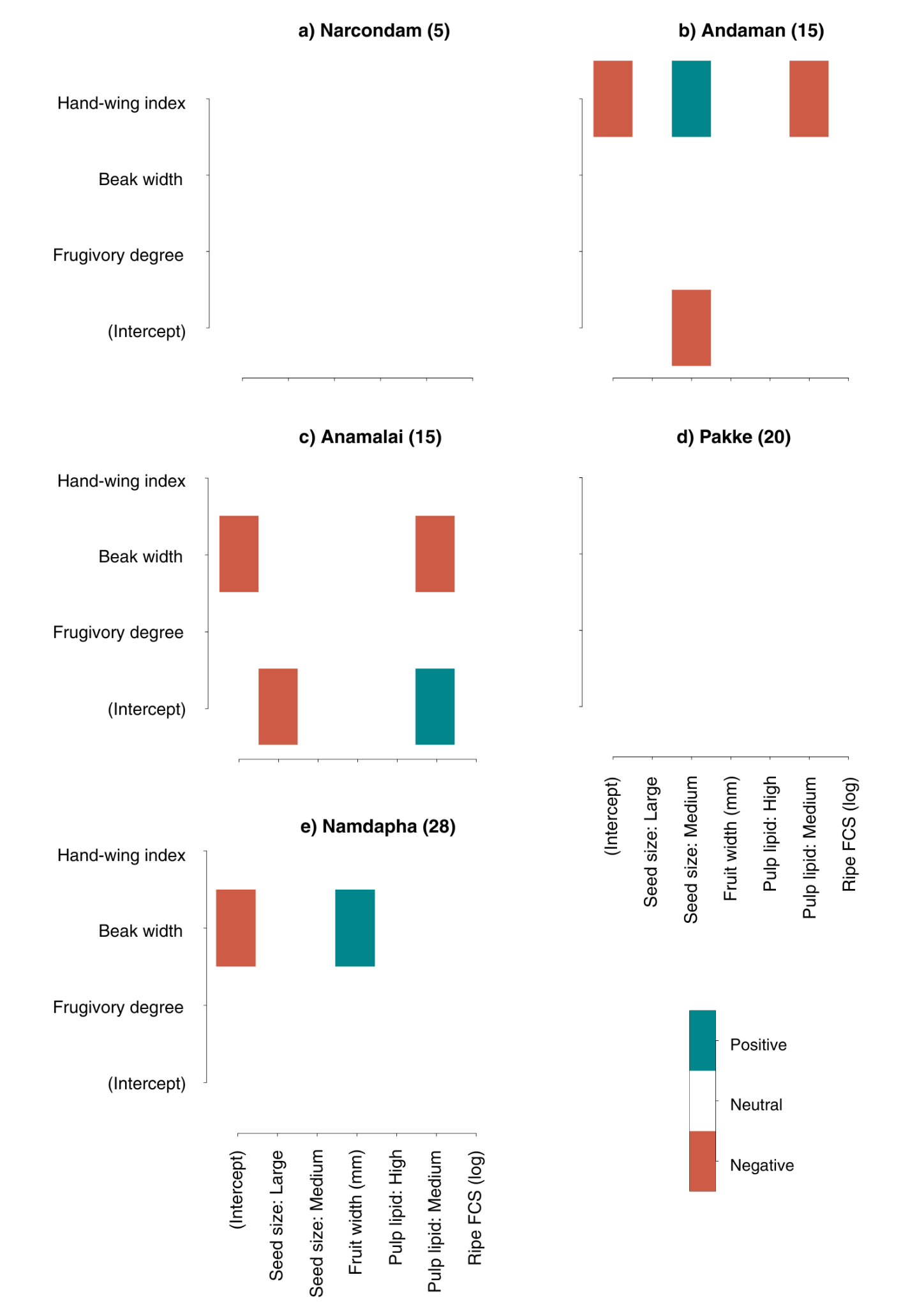

**Fig. S2.** Heatmaps showing the relationships between frugivore functional traits, namely, hand-wing index, beak width, and frugivory degree and fixed effects incorporated in the HMSC analysis, namely, seed size (small, medium, and large), fruit width (mm), pulp lipid (low, medium, and high), and log-transformed ripe fruit crop size (FCS). Each panel represent one study site, and the numbers in parentheses correspond to the number of frugivore species used in the analysis for that particular study site. The orange colour represents a positive relationship, turquoise represents a negative relationship, and white represents no relationship at a 95% posterior support level.

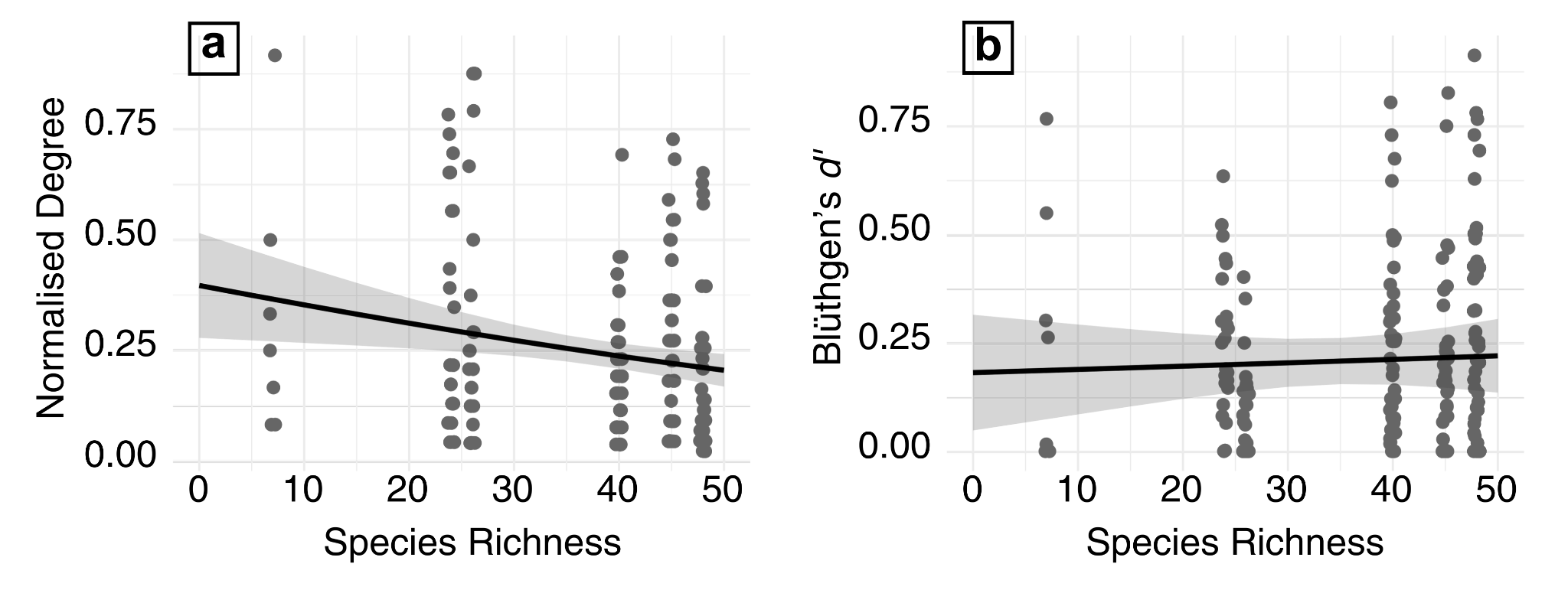

**Fig. S3.** Predicted relationship between the specialisation indices used in our study, namely, Normalised Degree or ND (a) and Blüthgen’s *d′* (b) and the site-level frugivore species richness (x-axis). Each dot represents the specialisation metric value (ND or *d′*) for an avian frugivore species at a particular site. The black line in each panel represents the best-fit line, and the shaded area the 95% confidence interval.
